# Subcutaneous vaccination with a live attenuated *Yersinia pseudotuberculosis* plague vaccine

**DOI:** 10.1101/822221

**Authors:** Anne Derbise, Chloé Guillas, Christiane Gerke, Elisabeth Carniel, Javier Pizarro-Cerdà, Christian E. Demeure

**Affiliations:** Yersinia Research Unit, Institut Pasteur, 28 rue du Dr Roux, 75724 Paris, France; Vaccine Programs, Institut Pasteur, 28 rue du Dr Roux, 75724 Paris, France

**Keywords:** bubonic plague, pneumonic plague, live vaccine, *Yersinia pestis*, *Yersinia pseudotuberculosis*, subcutaneous

## Abstract

A single oral inoculation to mice of the live attenuated *Yersinia pseudotuberculosis* VTnF1 strain producing an F1 pseudocapsule protects against bubonic and pneumonic plague. However oral vaccination can fail in humans exposed to frequent intestinal infections. We evaluated in mice the efficacy of subcutaneous vaccine injection as an alternative way to induce protective immunity, while reducing the dose and avoiding strain release in nature. A single subcutaneous dose of up to 10^8^ CFU induced dose-dependent antibody production. At the dose of 10^7^ CFU, i.e. 10 times less than via the oral route, it caused a modest skin reaction and protected 100% against bubonic and 80% against pneumonic plague, caused by high doses of *Yersinia pestis*. Bacteria migrating to lymph nodes and spleen, but not feces, were rapidly eliminated. Thus, subcutaneous injection of VTnF1 would represent a good alternative when dissemination in nature and human intestinal responsiveness are limitations.

## Introduction

Despite efficient antibiotic treatments, plague lethality in endemic regions remains superior to 10% [1], mainly due to the fast development of pathogenesis. Plague is endemic in more than 25 countries worldwide (including Madagascar, RDC, USA, China, Peru) and in addition to permanent foci, plague is reappearing since few decades and is categorized by WHO (World Health Organization) as a re-emerging disease [1]. No vaccine is widely available. A live attenuated *Y. pestis* vaccine is used in Russia and China but requires yearly boosting [2]. Two candidate recombinant vaccines under development in the USA and China are composed of F1 and LcrV (V) antigens. They have passed phase II clinical development but are not yet commercialized [3, 4].

We recently proposed to use a live *Y. pseudotuberculosis* to vaccinate against plague. This species originated from the same ancestor as the plague agent and is genetically very closely related but much less pathogenic. To this aim, a genetically defined *Y. pseudotuberculosis* strain was strongly attenuated by irreversible deletion of three loci encoding essential virulence factors (psaA, HPI, and yopK). It was modified to produce the *Y. pestis* F1 pseudocapsule by insertion of the *Y. pestis caf* operon, and named VTnF1. A single oral dose of VTnF1 generates both humoral and cell-mediated immune responses against multiple antigens of the bacteria and confers high-level protection against bubonic and pneumonic forms of plague [5].

VTnF1 was devised for the oral route because Yersiniae are enterobacteria which enter host gut secondary lymphoid tissues, strongly stimulate the host immune system, thus causing their own elimination. Although efficient for immunization, there are challenges for implementation. First, the oral route is not equally efficacious in different populations. Multiple intestinal infections alter the gut wall and immune system and reduce the efficacy of oral vaccination [6, 7] and most of the human population at risk for plague (Madagascar, Congo, Tanzania, etc.) is also exposed to such intestinal infections. Second, the dissemination of a live organism vaccine in the environment via feces may be considered unacceptable by those who fear GMOs and consider possible the reversion to more virulence [8, 9].

Vaccination by subcutaneous injection (sc; or hypodermal), intradermal or intramuscular injection could avoid these difficulties. The skin is an attractive site from an immunologic perspective: it contains many resident dendritic cells which capture antigens and migrate to lymph nodes to present antigens and recruit T lymphocytes [10]. Vaccination via skin generally requires less vaccine as compared to the oral route, in which a large part is eliminated by the intestinal transit. Bacteria injected sc also should not be disseminated in nature. We here examined the possibility to subcutaneously vaccinate mice against plague using the VTnF1 live attenuated candidate vaccine.

## Materials and methods

### Ethics statement

The Institut Pasteur animal facility is accredited by the French Ministry of Agriculture (B 75 15-01, issued on May 22, 2008), in compliance with French and European regulations (EC Directive 86/609, French Law 2001-486; June 6, 2001). The research protocol was approved by the French Ministry of Research (N° 2013-0038).

### Bacterial strains and culture conditions

The *Y. pseudotuberculosis* and *Y. pestis* isolates and their growth conditions were previously described [5, 11]. All experiments involving *Y. pestis* strains were performed in a BSL3 laboratory or level 3 animal room.

### Animal vaccination and *in vivo* analyses

Seven-week-old OF1 female mice (Charles River) were housed and manipulated in a BSL3 animal facility as described previously [11]. Vaccination consisted of subcutaneous (sc) injection of a VTnF1 suspension (10^5^, 10^6^, 10^7^, or 10^8^ CFU in 50μl saline) in the skin of the back. Mice were monitored daily for signs of prostration and aspect (ruffled hair) and were weighed. The skin reaction was scored as follows. 0: no papule; 1: not visible but perceptible by pinching the skin between fingers (≤ 1mm), 2: visible but small (1-2 mm) between fingers; 3: ≥ 2 mm between fingers. To examine *in vivo* VTnF1 dissemination, groups of mice were euthanized after predefined times and skin at the injection site (2 cm^2^), draining lymph nodes, and spleen were collected aseptically. They were homogenized in sterile PBS using 3 mm glass beads (soft tissues) or steel beads (skin) and an electric mill (TissueLyser®, Qiagen). Samples were plated on LB-agar containing Kanamycin and Spectinomycin. The presence of VTnF1 in feces was tested by plating 2-3 pellets per mouse.

### Plague challenge experiments

Mice were challenged four weeks after vaccination with *Y. pestis* resuspended in saline after growth at 28°C on LBH plates. Bubonic plague was induced by sc injection in the ventral skin. For pneumonic plague, it had been reported that too low volumes of bacteria instilled intranasally mostly stay in the upper respiratory tract [12, 13]. Using bioluminescent *Y. pestis* [5], we observed that lung colonization was obtained when 20 μl of bacteria (10 μl per nostril) were instilled (Supplementary Figure 1). Therefore, mice were infected as previously described [5], except the volume of 20 μl was used. Survival was followed for 21 days.

### Evaluation of the immune response

Mouse blood was collected three weeks after vaccination and serum IgG directed against F1 or a *Y. pestis* Δ*caf* sonicate were measured as previously described [5, 11].

### Statistical analyses

The Fisher exact test (for mice survival), the unpaired Mann-Whitney test (for bacteria numbers, animal weight, and antibody titers) were performed with Prism® (GraphPad).

## Results and discussion

### The VTnF1 candidate vaccine strain injected subcutaneously is avirulent and induces a protective immune response

Increasing doses of VTnF1 (10^5^ to 10^8^ CFU tested altogether in 24, 24, 46, and 38 mice respectively) were injected subcutaneously to OF1 mice in the back. No lethality and no clinical signs of illness were observed over a 4 weeks post vaccination follow-up, showing that VTnF1, strongly attenuated via the oral route [5], is also highly attenuated via a parenteral route (sc). This also indicates that the attenuation affected the bacteria’s ability to survive in the host body, and not only its capacity to infect the gut.

To evaluate protection against bubonic plague, mice received a single injection of VTnF1 (10^5^ to 10^8^ CFU) and were challenged 4 weeks after by ventral sc injection of *Y. pestis* (severe bubonic plague challenge: 10,000×LD_50_). High-level protection (96-100%) was obtained with the 10^7^ CFU dose and higher (Table 1). Therefore, protection via the sc route required a dose at least 10-times lower than via the oral route [5].

**Table 1:** Protection of mice by subcutaneous or intragastric vaccination with VTnF1 against bubonic and pneumonic plague. Mice having received a single subcutaneous dose of VTnF1 were challenged 4 weeks later by infection with *Y. pestis* CO92 s.c. (bubonic plague; 10^5^ CFU) or i.n. (pneumonic plague; 10^7^ CFU). Survival was recorded daily for 21 days. Survival from bubonic plague observed with VTnF1 given ig is reported here for comparison, and was first reported in Derbise A. *et al.* PLoS NTD. 2015. Protection significance compared to naïve mice was tested using Fisher’s two-tailed exact test: ***: p<0.001; ****: p<0.0001

| Vaccine dose<br>(CFU) | Subcutaneous VTnF1 |  |  |  |  |  |  |  | Intragastric VTnF1 |  |  |  |  |
| --- | --- | --- | --- | --- | --- | --- | --- | --- | --- | --- | --- | --- | --- |
|  | Bubonic plague |  |  |  |  | Pneumonic plague |  |  | Bubonic pl. |  |  | Pneumonic pl. |  |
|  | 0 | 10<sup>5</sup> | 10<sup>6</sup> | 10<sup>7</sup> | 10<sup>8</sup> | 0 | 10<sup>7</sup> | 10<sup>8</sup> | 0 | 10<sup>7</sup> | 10<sup>8</sup> | 0 | 10<sup>8</sup> |
| Survivors / total | 0/16 | 13/24 | 23/24 | 16/16 | 8/8 | 0/23 | 24/30 | 25/30 | 1/16 | 6/8 | 13/14 | 0/8 | 13/15 |
| % survival | 0 | 54 | 96 | 100 | 100 | 0 | 80 | 83 | 6 | 75 | 93 | 0 | 87 |
| Significance |  | *** | **** | **** | **** |  | **** | **** |  | * | *** |  | **** |

The protection induced by sc and ig vaccination against pneumonic plague were compared using recently modified conditions to induce airways infection modified recently (see materials and methods). Both routes conferred around 80% protection (Table 1) against pneumonic plague caused by a high dose of *Y. pestis* (10^7^ CFU = 3,300 × LD_50_) and increasing the sc dose to 10^8^ CFU did not increase protection.

*Y. pestis* specific IgG were measured in the serum of vaccinated mice. Vaccination induced a strongly dose-dependent production of IgG directed to either F1 (Fig.1A) or all other *Yersinia* antigens (F1- *Y. pestis*; Fig. 1B). Antibody titers were compared to those of oral VTnF1. Sc injection induced comparable amounts of IgG against F1, and more IgG against other *Yersinia* antigens (F1- *Y. pestis*; Fig. 1A & B) than ig inoculation. VTnF1 at dose 10^7^ given ig yields heterogenous IgG titers and lower protection (Fig 1 and [14]). Altogether, the vaccine given sc is as efficient in terms of antibody production (Fig. 1) and protection (Table 1), as when it is given ig at a higher dose. One explanation to the higher IgG response after sc vaccination may be that bacteria trapped in skin are available to trigger the immune system, whereas part of bacteria via ig is killed by acidity in the stomach, or fail to bind PP and are washed away in feces. Similarly, Zhao and co-workers recently reported that the sc route provided higher protection than the oral route for mice vaccination with a live *Salmonella* vector producing *Salmonella* or *Bordetella* antigens [15].

**Figure 1:**
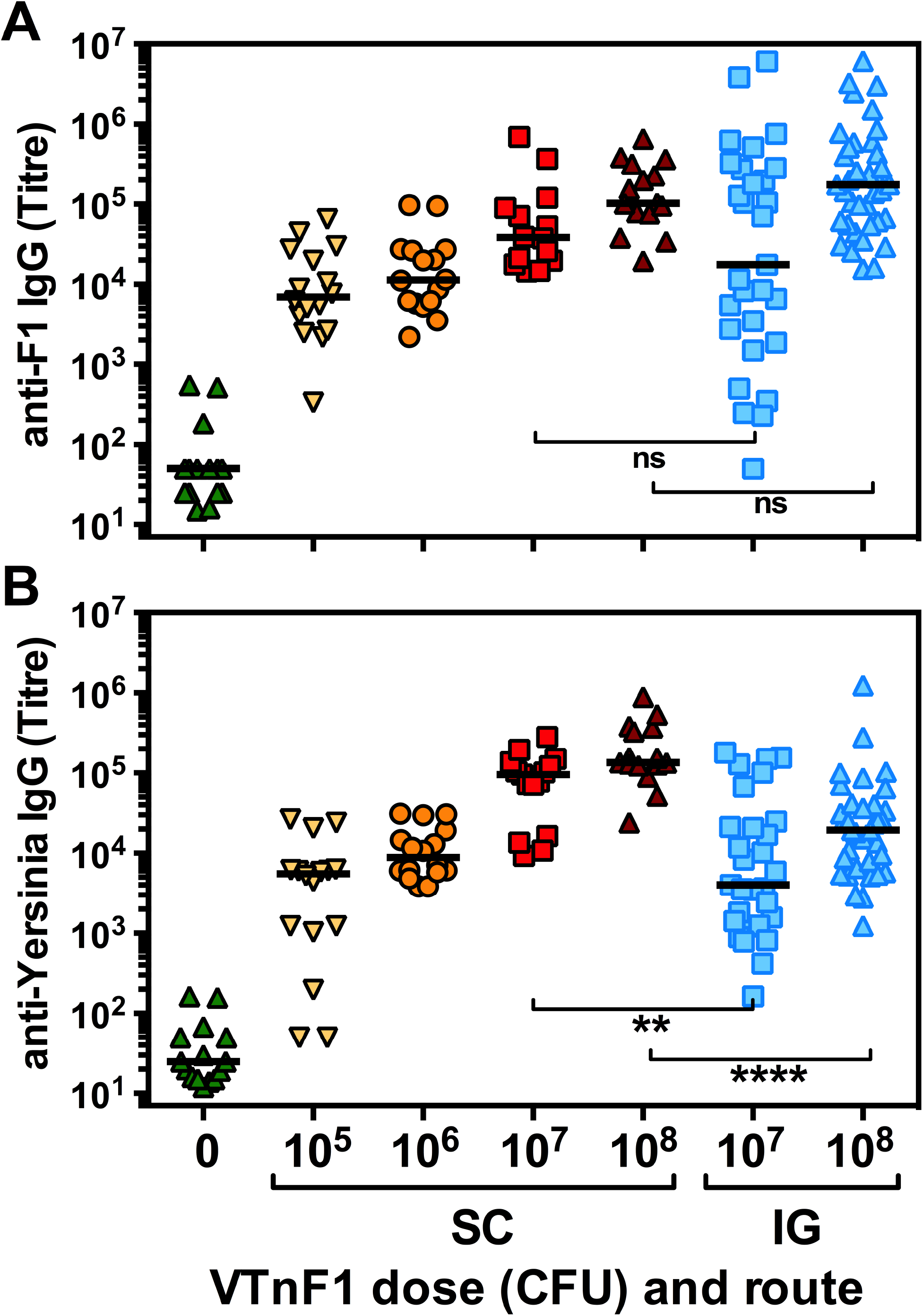
Immune response of vaccinated mice. Groups of mice were injected sc (14 /group gathered from 2 experiments) or inoculated ig (30-37 mice gathered from 3 experiments) with graded doses of live VTnF1 and blood was taken 3 weeks. Serum IgG against purified F1 (A) or other *Y. pestis* antigens (CO92 Δ*caf* lysate: all antigens except F1; B) were measured in sera by Elisa. Mann-Whitney test; *: p≤0.05; ***: p≤0.001; ****: p≤0.0001; ns: not significant.

A key reason to evaluate the cutaneous route was that the gut immune system of subjects exposed to repeated oral infections has a reduced ability to respond to immunization, thus reducing the efficacy of oral vaccines [6, 7]. Such an impairment has so far not been reported regarding the skin immune system, suggesting that skin vaccination remains a valid prophylactic approach in these populations.

### VTnF1 causes little side effects and disseminates transiently in vivo

Although the subcutaneous route is widely used for live attenuated viruses (Varicella, measles, mumps [16]), injection of live attenuated bacteria is not usual, possibly due to side effects with candidate vaccines tested previously [17]. Because VTnF1 is highly attenuated and is efficient at the dose of 10^7^ CFU, we characterized side-effects at that dose. After injection, half of mice had ruffled fur at day 2, and most had recovered at day 5. Skin papules at the site of injection were always palpable (Fig. 2A), but were generally not visible by eye, except in 10% of mice who exhibited a local hair loss, which always regrew to normal afterwards. A reduction of weight acquisition was observed at day 3, but growth rapidly normalized later on (Fig. 2B). Because humans are more sensitive to LPS than mice, a reaction of human tissues to LPS contained in the vaccine may happen, and this will carefully be examined in future developments. LPS detoxification by deletion of *msbB* or htrB genes controlling lipid-A acylation could be considered, as previously reported in *Shigella* [18].

**Figure 2:**
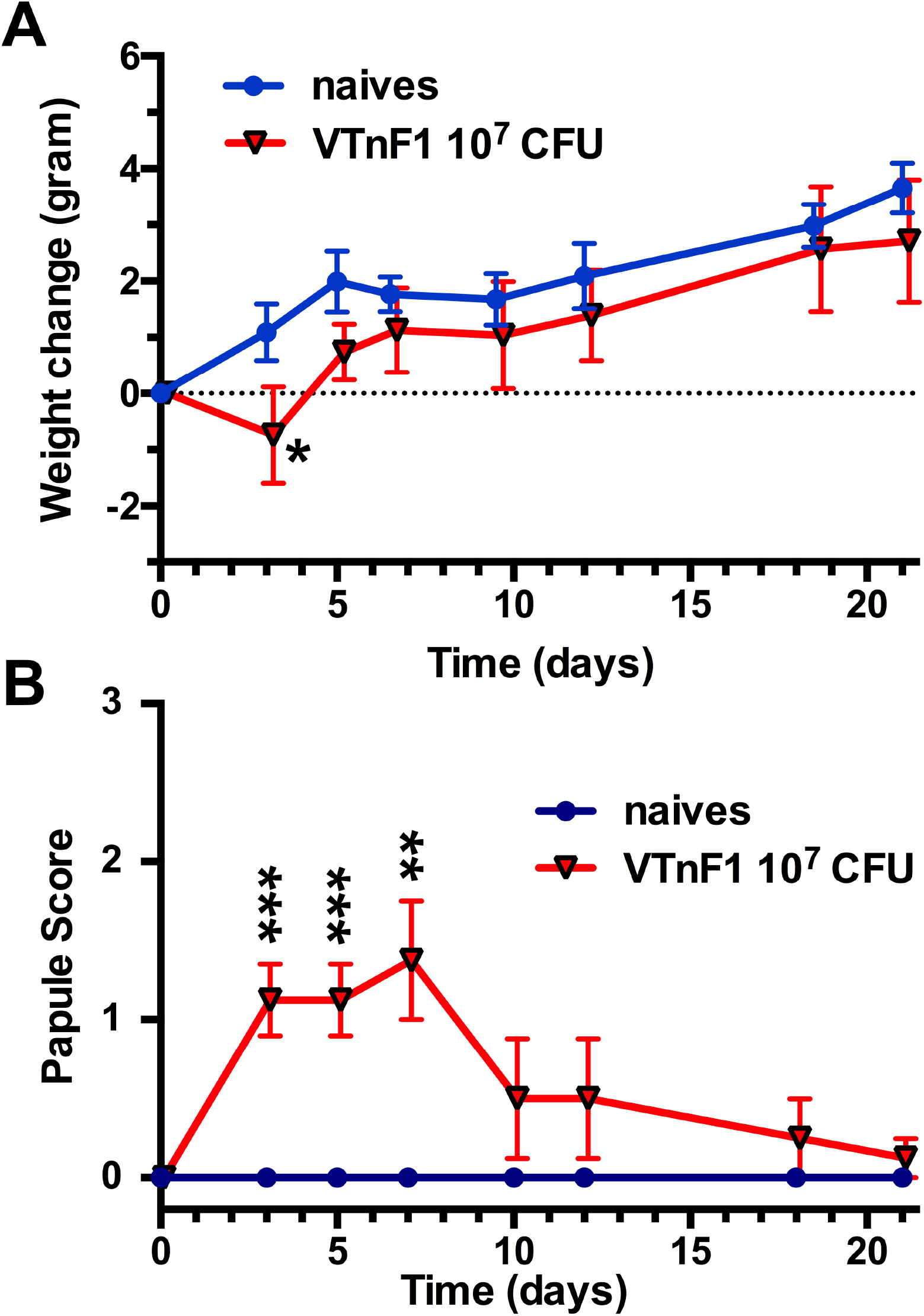
Follow-up of vaccinated mice. Mice were vaccinated sc with VTnF1 (triangles), or were left unvaccinated (naïves: circles) and were followed for 3 weeks for their weight (A) and the size of the reaction at the skin site of vaccination (B; 0: undetectable; 1: not visible but perceptible (≤ 1mm) by pinching, 2: visible but small (1-2 mm); 3: ≥ 2 mm). Shown are means ± s.e.m. of 15 (A) or 8 (B) mice per group. The statistical significance was evaluated using the unpaired Student t test). *: p≤0.05; **: p≤0.01; ***: p≤0.001.

VTnF1 dissemination from the skin to the draining lymph nodes (LN) and the spleen was examined (Fig. 3). Bacteria persisted at the skin entry site up to day 5 and then declined, so that most mice were negative after 3 weeks. At day 1-2, VTnF1 was observed in spleen and draining lymph nodes, suggesting a simultaneous migration to spleen via blood and to lymph nodes via lymphatics. VTnF1 loads peaked in all sites at day 5, prior to a progressive disappearance in LN and spleen after day 9. This transient persistence was therefore comparable, in its evolution and levels, to that observed after oral vaccination with the higher dose, in good agreement with the comparable immunization.

**Figure 3:**
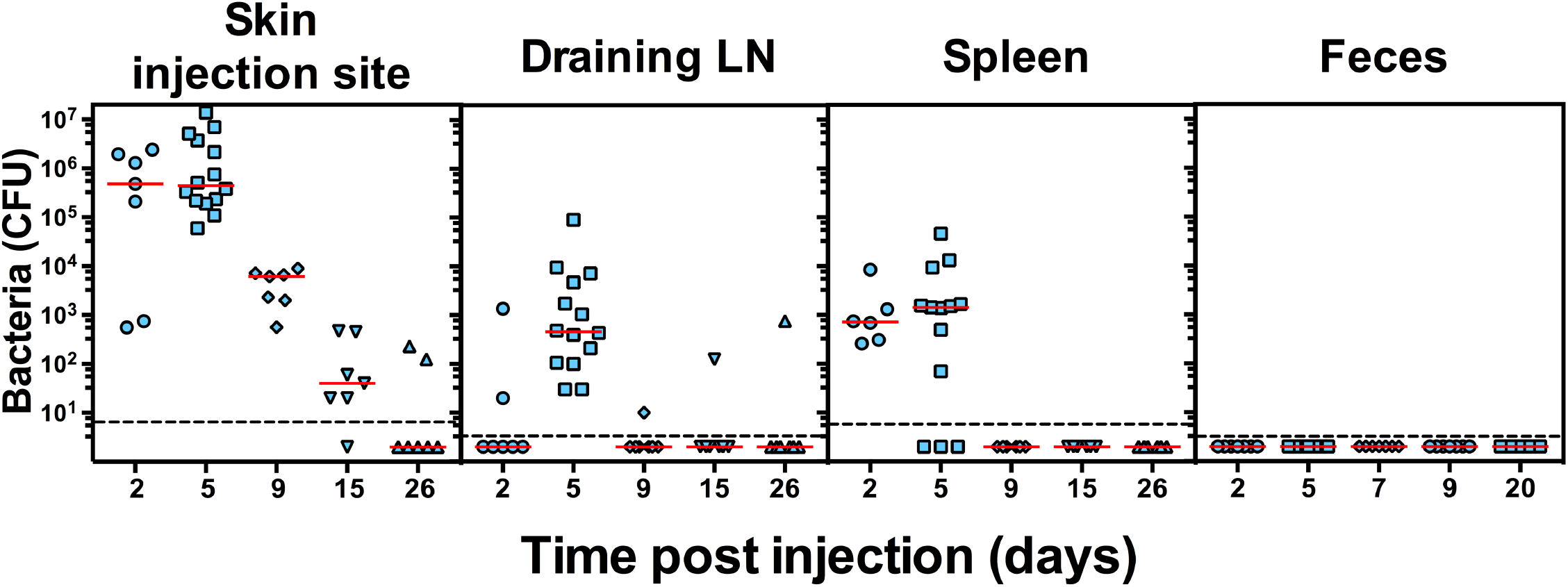
In vivo dissemination of sc-injected VTnF1. Groups of mice were injected sc with 10^7^ CFU VTnF1 and were sacrificed at predetermined times. Organs and feces were collected and homogenized for bacteria load determination. Individual results from 7 mice per group are shown for skin (2 cm^2^ around the site of injection), DLN from both sides, the whole spleen and 2-3 fecal pellets. The median is shown as a red horizontal bar. The dotted line indicates the limit of detection.

The subcutaneous tissue in which the bacteria were injected, also named hypodermis, is a well vascularized layer of adipose tissue and the most used skin layer for delivery of live attenuated virus vaccines. Although it is less drained by lymphatics than the actively surveyed dermis layer, injected particles succeed to reach LN via lymph to trigger adaptive immunity [19]. Because VTnF1 rapidly reached both LN and spleen (day 1), it must have gained access to both blood and lymphatic vessels, and this may result from a mechanical damage caused by injection. Alternatively, antigens could also be carried to LN by specialized antigen presenting cells. Because the hypodermis is naturally devoid of resident immune cells, these would be recruited from dermis or blood.

The risk of bacteria passage in the environment was evaluated by searching for VTnF1 in feces. Feces were analyzed regularly up to one month post inoculation, and no VTnF1 was detected (Fig. 3). Although it seems trivial that skin is not directly connected to the intestine, the transit of bacteria via blood could have redirected it to unexpected organs, as was observed for *Listeria monocytogenes* moving from blood to the intestine [20]. The absence of vaccine in feces demonstrates that it did not reach the gut lumen, and that fecal dissemination of this GMO in nature is very unlikely. For future use, we are re-constructing the VTnF1 strain without any antibiotics-resistance cassettes (unpublished data), and this will increase safety for both vaccinated subjects and in case of accidental release.

In conclusion, we have shown that subcutaneous vaccination with the recombinant attenuated *Y. pseudotuberculosis* vaccine strain VTNF1 efficiently immunized and protected mice against bubonic and pneumonic plague, with an optimal dose lower than required for oral vaccination. Because it also avoids dissemination of the vaccine in nature, the hypodermal route may be an interesting alternative to oral vaccination when its efficacy is impaired, such as often observed in developing countries.

## Acknowledgements

The project was supported by an ANR Emergence grant (ANR-12-EMMA-0011-01) and an Institut Pasteur Accelerating Preclinical Candidates - GPF– Vaccinology 2015 grant (GPFVacc2-08). The authors would like to thank the Institut Pasteur Central animal facility team for their support with mouse hosting and care.

---

**Figure S1:**
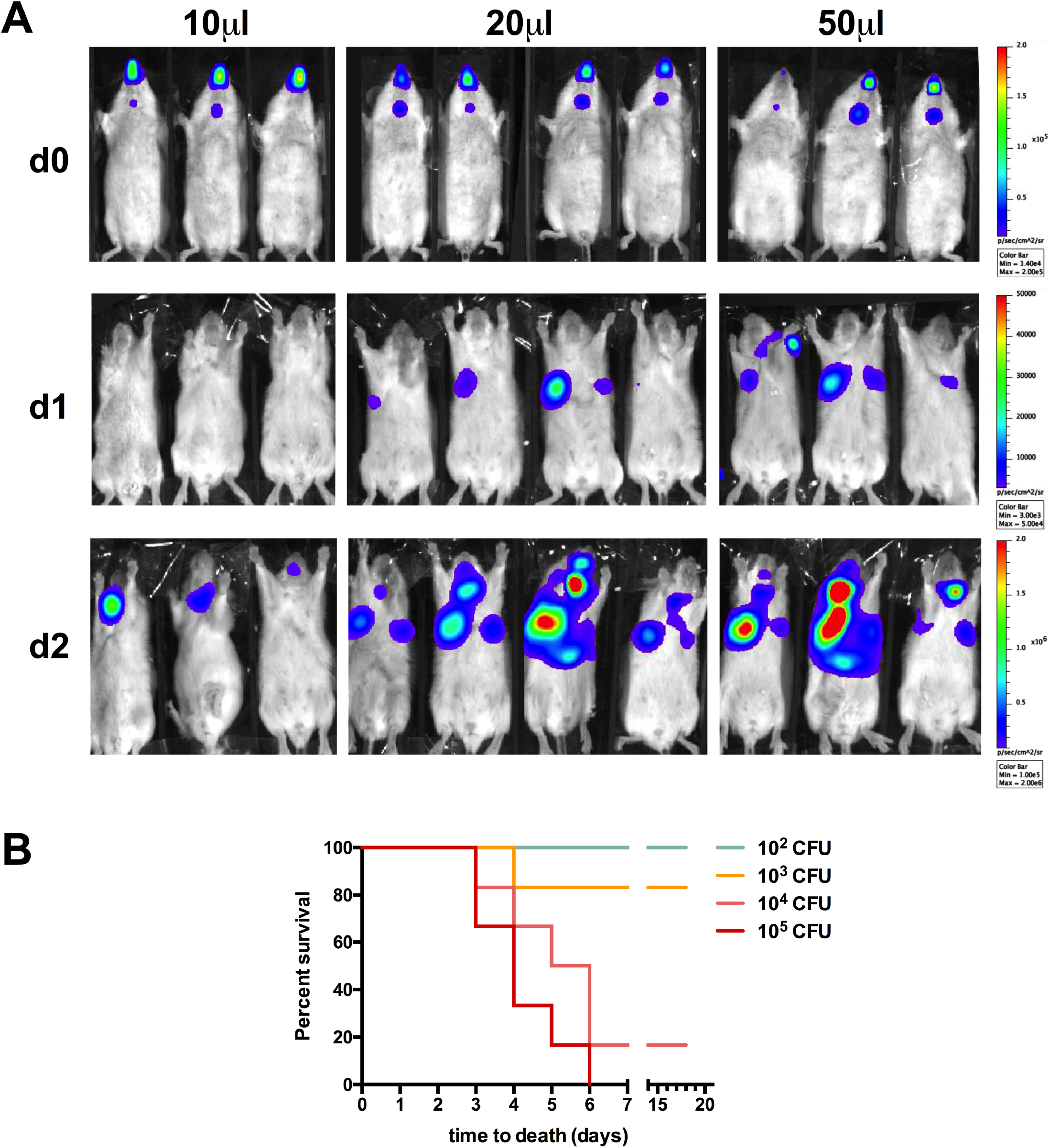
Influence of the volume instilled intranasally on lung infection with *Y. pestis* as a model of pneumonic plague. (A) Mice anesthesized using Xylazine + Kétamine and laying on the back were infected by intranasal instillation of 10^7^ CFU of the bioluminescent *Y. pestis* CO92::Tn7-*Pail-lux* (5). To this aim, 10, 20 or 50 microliters were progressively presented as small drops in front of nostrils for inhalation (half dose in each nostril). Whole body bioluminescence of the animals was recorded at regular time intervals using an IVIS® camera. Shown are white light photographs merged with the bioluminescence signal, scaled in the right margin. (B) The median lethal dose (LD_50_) of *Y. pestis* instilled in nostrils as 20 microliters was calculated by infecting groups of mice with increasing doses of *Y. pestis*, and animal death from infection was followed for 21 days. The LD_50_ calculated using the Spearman-Karber method was found equal to 3160 CFU (95% confidence interval: 1174–8517).

